# U1 snRNA blockade regulates DNA repair genes, DNA damage, and cisplatin sensitivity of lung cancer cells

**DOI:** 10.64898/2026.08.27.747528

**Authors:** Alexandre Devaux, Céline M. Labbe, Stéphan Vagner, Martin Dutertre

**Affiliations:** Institut Curie, Université PSL, CNRS UMR3348, INSERM U1368, 91400 Orsay, France; Université Paris-Saclay, CNRS UMR3348, INSERM U1368, 91400 Orsay, France; Equipe Labellisée Ligue Nationale Contre le Cancer

**Keywords:** Intronic polyadenylation, U1 snRNA, Antisense oligonucleotide, Genotoxic agent, Lung cancer

## Abstract

Previous studies revealed a crosstalk between intronic polyadenylation (IPA) and the DNA damage response (DDR). Indeed, genotoxic agents, including radiations and anticancer drugs (*e.g.*, cisplatin that crosslinks DNA), regulate the ratio of IPA to last-exon transcripts in many genes. Conversely, multiple genes involved in the DDR, especially homologous recombination, are regulated at the IPA level. The U1 small nuclear RNA (snRNA) widely represses IPA, thereby enhancing full-length gene transcription. However, besides its implication in IPA regulation by ultraviolet-C radiation, little is known about U1 snRNA effects on the DDR and on cell sensitivity to genotoxic agents. Here, we show that U1 snRNA blockade using an antisense oligonucleotide (U1-AMO) in lung cancer cell lines enhances cell growth inhibition by cisplatin, through an increase in cisplatin-induced DNA damage. 3’-seq analysis indicates that U1 snRNA blockade represses full-length mRNA expression of multiple genes of the nucleotide-excision repair and Fanconi anemia pathways, which are involved in the repair of cisplatin-DNA crosslinks. Our 3’-seq analyses also reveal that moderate doses of U1-AMO and cisplatin upregulate the IPA:LE isoform ratio in overlapping but distinct sets of genes, and that U1-AMO prevents cisplatin effects on the IPA:LE ratio in a large subset of genes. Altogether, these data extend the crosstalk between IPA and the DDR and suggest that U1 snRNA targeting may be used to sensitize cancer cells to genotoxic agents.

## INTRODUCTION

The U1 small nuclear RNA (snRNA), as part of the U1 small nuclear ribonucleoprotein (snRNP), is a core component of the spliceosome. U1 snRNA hybridization with pre-messenger RNA (pre-mRNA) 5’ splice sites mediates their recognition, that is required for splicing. The U1 snRNP also has splicing-independent activities, including stimulation of transcription initiation (Kwek et al. 2002; Damgaard et al. 2008) and elongation (Mimoso and Adelman 2023). In addition, U1 snRNA binding to pre-mRNA can impact the use of a nearby cleavage and polyadenylation site (polyA site) (Gunderson et al. 1998; Vagner et al. 2000). In fact, the main RNA effect observed upon U1 snRNA blockade using an antisense morpholino oligonucleotide (U1-AMO) targeting its 5’ splice site-binding region is an increase in the use of intronic polyadenylation (IPA) sites (Kaida et al. 2010; Berg et al. 2012; Vorlová et al. 2011; Andersen et al. 2012; Oh et al. 2017). IPA sites are sometimes referred to as premature cleavage and polyadenylation (PCPA) sites, although a subset of them are used to generate transcript isoforms with alternative last exons. As a result of PCPA, U1-AMO decreases the synthesis of full-length mRNAs in thousands of genes, as shown by RNA-seq analysis of nascent RNA (Oh et al. 2017).

Genotoxic agents, including both radiations and drugs, can be used to kill cancer cells through the induction of DNA lesions. The so-called DNA damage response (DDR) comprises the recognition of various types of DNA lesions, their repair by multiple pathways, the activation of cell-cycle checkpoints, and other processes, leading eventually to either cell survival or cell death (Ciccia and Elledge 2010; Dutertre et al. 2014b). Among genotoxic anticancer agents is cisplatin, which is widely used in several cancer types such as lung cancer. Cisplatin induces both intra– and inter-strand DNA crosslinks, that are repaired mainly by the nucleotide-excision repair (NER) and Fanconi anemia (FA) pathways, respectively (Rocha et al. 2018).

A reciprocal crosstalk between IPA and the DDR has emerged in recent years (Dutertre et al. 2021). On the one hand, the IPA to last-exon (LE) isoform ratio is widely regulated by genotoxic agents (Dutertre et al. 2014a; Williamson et al. 2017; Devany et al. 2016; Chakraborty et al. 2022; Devaux et al. 2025), with mainly upregulation in the case of ultraviolet-C irradiation (Devany et al. 2016; Williamson et al. 2017) and cisplatin treatment (Devaux et al. 2025). For both of these agents, IPA:LE regulation events are enriched in long genes and are mediated at least in part by a repression of transcription processivity (Williamson et al. 2017; Devaux et al. 2025). In addition, following ultraviolet-C irradiation, U1 snRNA levels are decreased, and its overexpression prevents several IPA:LE regulation events and apoptosis induction (Devany et al. 2016).

On the other hand, genes that are regulated at the level of IPA are enriched in DDR genes. Indeed, depletion or inhibition of cyclin-dependent kinase 12 (CDK12) increases the IPA:LE isoform ratio and decreases transcription processivity in multiple genes involved in the DDR –especially in homologous recombination– and sensitizes cells to genotoxic agents, presumably due to the IPA-mediated regulation of DDR genes (Chirackal Manavalan et al. 2019; Dubbury et al. 2018; Fan et al. 2020; Krajewska et al. 2019; Quereda et al. 2019; Tien et al. 2017). However, CDK12 interacts with and phosphorylates many proteins in addition to the RNA polymerase II large subunit (Krajewska et al. 2019; Tien et al. 2017; Bartkowiak et al. 2019) and, through the phosphorylation of a translation factor, has been involved in translational regulation of a DDR protein (CHK1) and of multiple mitosis proteins controlling mitotic chromosome stability (Choi et al. 2019).

In contrast with CDK12, the effects of the U1 snRNA on DNA damage and cell sensitivity to genotoxic and/or anticancer agents have been little studied and were therefore the subject of the present study. For this, because lung cancer is the leading cause of cancer-related death worldwide and is widely treated with platinum compounds in the clinic (Ashrafi et al. 2022), we studied the sensitivity of lung cancer cell lines to cisplatin.

## RESULTS

### U1 snRNA blockade enhances cell growth inhibition by cisplatin in lung cancer cells

Because U1 snRNA widely represses IPA, which is known to be enriched in DDR genes, we tested the ability of U1-AMO to sensitize cancer cell lines to a genotoxic agent. For this, we used two cell lines (A549 and H358) derived from non-small cell lung cancer (the main type of lung cancer). Cells were transfected with either a control oligonucleotide (CTRL-AMO) or various doses of U1-AMO (complemented with CTRL-AMO to achieve the same total amount of AMO in every condition in a given cell line). Then, cells were treated or not with cisplatin and finally assayed for cell viability by WST1 assay. In the absence of cisplatin, U1-AMO decreased the growth of both cell lines, when compared to CTRL-AMO, with a nice dose-dependent effect in H358 cells (Fig. 1A, vehicle). This is consistent with previous studies in other cell lines.

**Fig. 1:**
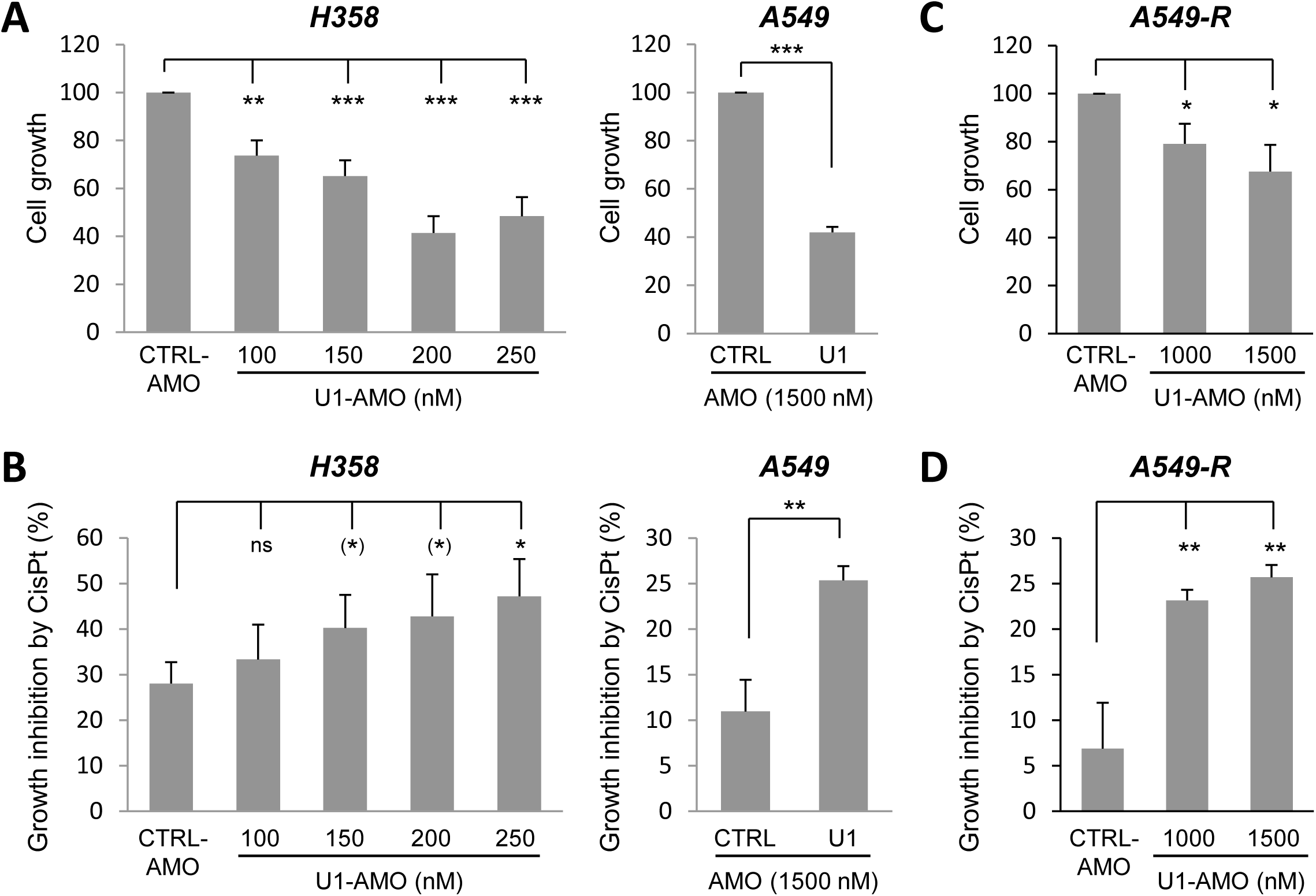
U1 snRNA blockade sensitizes cells to growth inhibition by cisplatin. **A-B**, H358, A549 and A549-R cells were electroporated with CTRL– and/or U1-AMO and subsequently treated with either cisplatin (CisPt at 10, 5 or 15 µM, respectively) or vehicle. In each panel, the total amount of AMO (CTRL + U1) is the same in every condition. Cell growth was measured by WST1 assay. **A**, cell growth in the absence of cisplatin. **B**, Cell growth inhibition by cisplatin, in cells with prior electroporation with either CTRL-AMO or U1-AMO.

To look at cell growth inhibition by cisplatin, we used cisplatin doses that had mild effects in CTRL-AMO electroporated cells, and for each dose of U1-AMO we compared cell viability with the cisplatin/U1– AMO combination *versus* U1-AMO alone. In both cell lines, when compared to CTRL-AMO, prior transfection with U1-AMO enhanced the inhibition of cell growth by subsequent cisplatin treatment, and there was a nice dose-dependent effect of U1-AMO in H358 cells (Fig. 1B). Thus, in both cell lines, U1 snRNA blockade sensitized cells to growth inhibition by cisplatin.

Finally, to test whether U1-AMO may also sensitize cells that acquired resistance to cisplatin, we used the A549-R cell line that was previously obtained by selection of A549 cells with increasing doses of cisplatin (Michels et al. 2013). As in parental A549 cells, U1-AMO decreased the growth of A549-R cells in the absence of cisplatin and increased cell growth inhibition by cisplatin (Fig. 1C-D).

### U1 snRNA blockade prevents the ability of cisplatin to inhibit cell cycle progression

Because genotoxic agents block cell cycle progression, and because little is known on U1-AMO effects on the cell cycle, we analyzed cell distribution in cell cycle phases following treatment with U1-AMO and cisplatin either alone or in combination, as above. For this, we carried out FACS analysis using propidium iodide and bromodeoxyuridine (BrdU) (Fig. 2A-B). In both cell lines, the G0-G1 to G2-M ratio, which reflects the ability of cells to progress from G2-M to G1 phase, was decreased by both U1-AMO and cisplatin alone (Fig. 2C-D, left panels), which is consistent with their growth-inhibitory effects.

**Fig. 2:**
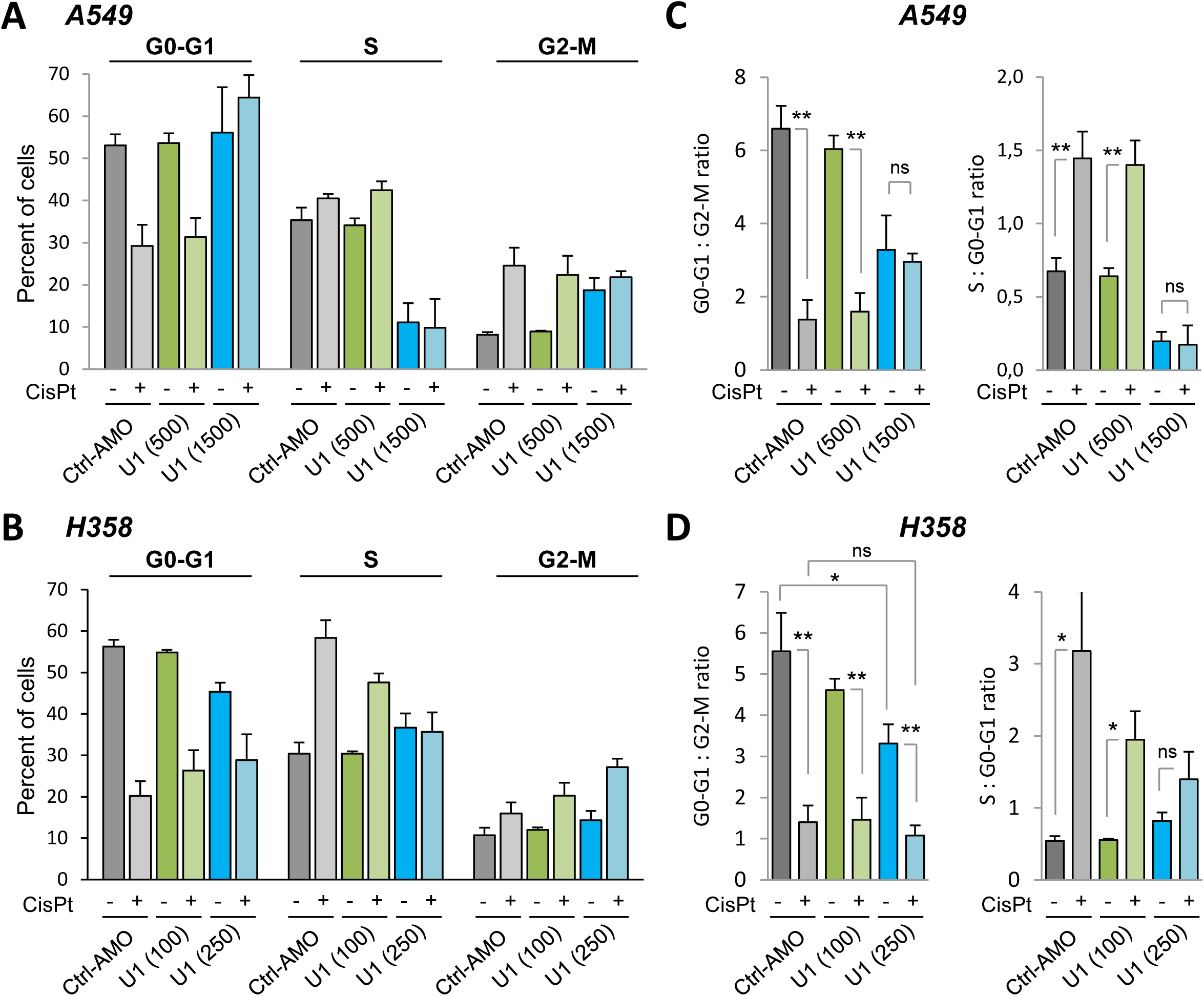
U1 snRNA blockade prevents the ability of cisplatin to inhibit cell cycle progression. H358 and A549 cells were electroporated with CTRL– and/or U1-AMO and subsequently treated with either cisplatin (at 10 or 5 µM, respectively) or vehicle. In each panel, the total amount of AMO (CTRL + U1) is the same in every condition. Cell cycle distribution was analyzed by FACS with propidium iodide and BrdU.

However, in both cell lines, U1-AMO at least partially prevented the effects of cisplatin on the G0-G1 to G2-M ratio (Fig. 2C-D, left panels; in H358 cells, the fold effect of cisplatin is decreased from 4.0 to 3.1). Regarding the S to G0-G1 ratio, it was increased by cisplatin in both cell lines, suggesting a replication blockade (Fig. 2C-D, right panels; it was also decreased by U1-AMO alone in A549 but not H358 cells). Again, U1-AMO prevented these cisplatin effects at least partially (Fig. 2C-D, right panels). Thus, the ability of U1 snRNA blockade to enhance the growth-inhibitory effect of cisplatin (Fig. 1B-C) is not due to an enhancement of the cell cycle effects of the drug. Instead, U1 snRNA blockade prevents (at least partially) the ability of cisplatin to inhibit cell cycle progression.

### U1 snRNA blockade increases cisplatin-induced DNA damage

Then, we tested whether cell sensitization to cisplatin by U1-AMO might be due to decreased repair of cisplatin-induced DNA damage. For this, we measured DNA damage by immunofluorescence of the γH2AX histone mark, a well-known marker of DNA damage (Rocha et al. 2018). In both A549 (Fig. 3A-B) and H358 cells (Fig. 3C-D), γH2AX staining (quantitated as either mean fluorescence intensity or foci number) was increased by cisplatin, as expected, while U1-AMO alone had little effect. Importantly, cisplatin-induced γH2AX staining was further enhanced by prior treatment with U1-AMO (in a dose-dependent manner). These data indicate that U1-AMO decreases the repair of cisplatin-induced DNA damage.

**Fig. 3:**
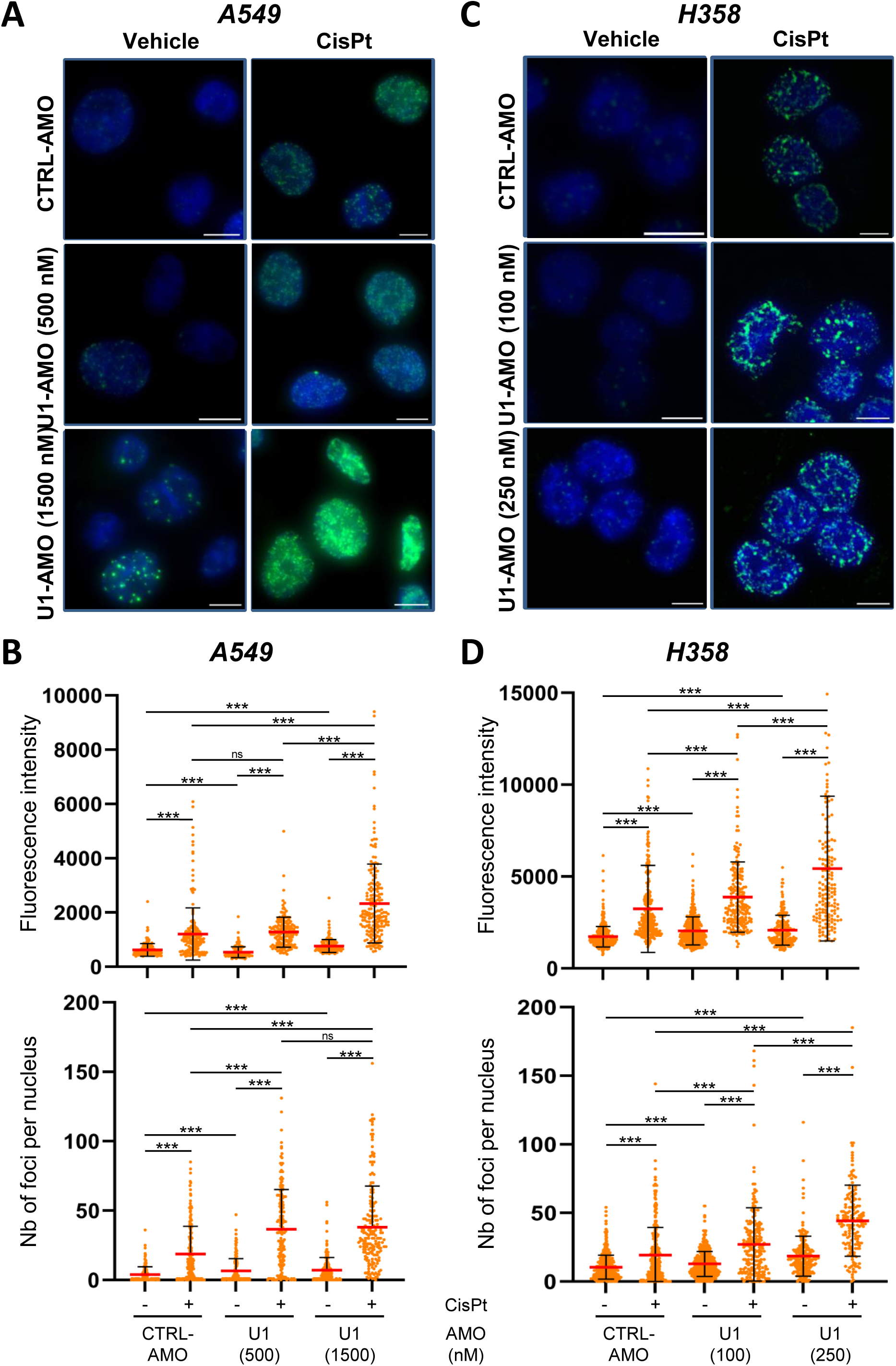
U1 snRNA blockade increases cisplatin-induced DNA damage. H358 and A549 cells were electroporated with CTRL– and/or U1-AMO and subsequently treated with either cisplatin (at 10 or 5 µM, respectively) or vehicle. In each cell line, the total amount of AMO (CTRL + U1) is the same in every condition. Immunofluorescence analysis of γH2AX. **A**, Typical images of γH2AX (green) and DAPI (blue) staining. Bars represent 10 µm. **B**, Quantification of γH2AX staining in a representative experiment. Each dot corresponds to a cell nucleus; the median and standard deviation are shown.

### U1 snRNA blockade represses genes of the NER and FA repair pathways

Then, to investigate how U1-AMO decreases the repair of cisplatin-induced DNA damage, we analyzed the impact of U1-AMO on gene expression globally. For this, we used 3’-seq (RNA-seq focused on the 3’-end of polyadenylated transcripts), because U1-AMO is known to increase PCPA, leading in turn to a decrease of transcription processivity towards the last exon in long genes (Oh et al. 2017). Consistently, treatment of A549 cells with U1-AMO at 500 and 1500 nM led to an increase of the IPA:LE isoform ratio in 1475 and 5776 genes, respectively (Fig. 4A, right panel, and Fig. S1A-B; these data will be presented in more detail in the next section). Among these genes, 22 and 535 had decreased expression of their last exon by at least 4-fold by U1-AMO at 500 and 1500 nM, respectively (Fig. 4A, right panel, and Supplemental Tables S1 and S2).

**Fig. 4:**
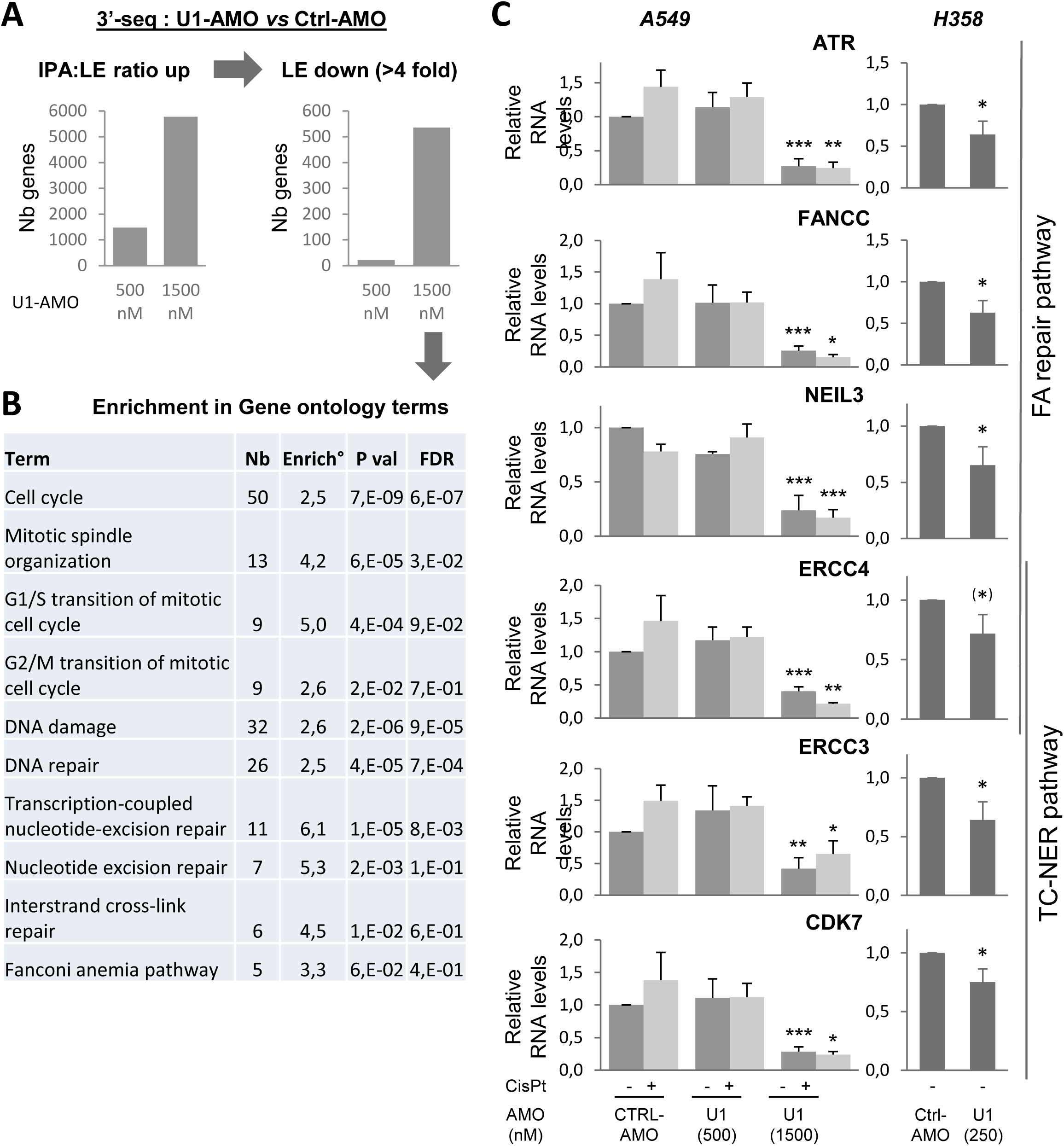
U1 snRNA blockade represses genes of the NER and FA repair pathways. A549 cells were electroporated with CTRL– and/or U1-AMO (with a total of 1500 nM AMO in every condition) and subsequently treated with either cisplatin at 5 µM or vehicle. **A**, 3’-seq analysis, comparing A549 cells electroporated with U1-AMO *versus* Ctrl-AMO. Number of genes with IPA:LE isoform ratio up-regulation (p<0,05, left panel) and last-exon level down-regulation (at least 4-fold, right panel). **B**, Enrichment analysis of gene ontology terms in the indicated list of genes. Only terms related to the DDR and cell cycle are shown. **C**, RT-qPCR analysis using primers located in the last exon of indicated genes in A549 and H358 cells electroporated with CTRL– and/or U1-AMO, as indicated.

The 535 repressed genes were enriched in genes involved in cell cycle (most notably, mitotic spindle organization and G1/S and G2/M transitions of mitotic cell cycle) and DNA repair (Fig. 4B and Supplemental Table S3). The most enriched DNA-repair pathways were the NER (especially, transcription-coupled [TC]-NER) and interstrand cross-link (especially FA) pathways (Fig. 4B). Remarkably, these are the two main repair pathways of cisplatin-DNA crosslinks. We validated by RT-qPCR that U1-AMO decreased in a dose-dependent manner the expression of the last exon of genes involved in the FA (*e.g.*, *ATR*, *FANCC*, *NEIL3*), TC-NER (*e.g.*, *ERCC3*, *CDK7*) or both pathways (*e.g.*, *ERCC4*) in A549 cells (Fig. 4C, left panels). Similar effects were seen with the combination of U1-AMO and cisplatin. Finally, these genes were also repressed by U1-AMO in H358 cells (Fig. 4C, right panels). Altogether, these data indicate that U1 snRNA blockade decreases the expression of the full-length mRNAs of multiple genes involved in the FA and TC-NER pathways of DNA repair.

### U1 snRNA blockade partially prevents cisplatin effect on the IPA:LE isoform ratio

As both U1-AMO ((Oh et al. 2017) and see above) and cisplatin (Devaux et al. 2025) were shown to regulate IPA isoforms in many genes, our 3’-seq analyses in A549 cells also allowed us to compare their effects, alone and in combination, on the IPA:LE isoform ratio on a genome-wide scale. As expected, when compared to control cells that received CTRL-AMO and vehicle, both U1-AMO and cisplatin agents used alone regulated IPA:LE events in many genes, with mainly upregulation events (Fig. 5A-B, lanes 1, 2 and 6). U1-AMO upregulated IPA:LE events in a dose-dependent manner, with 2088 events in 1475 genes at 500 nM, and 29677 events in 5776 genes at 1500 nM (Fig. 5A-B, lanes 2 and 6; and Supplemental Fig. S1A-B). Meanwhile, treatment with cisplatin alone at 5 µM upregulated 2881 IPA:LE events in 1396 genes, when compared to control cells (Fig. 5A-B lane 1, Supplemental Fig. S1E and Supplemental Table S4). Going further, we made three interesting observations.

**Fig. 5:**
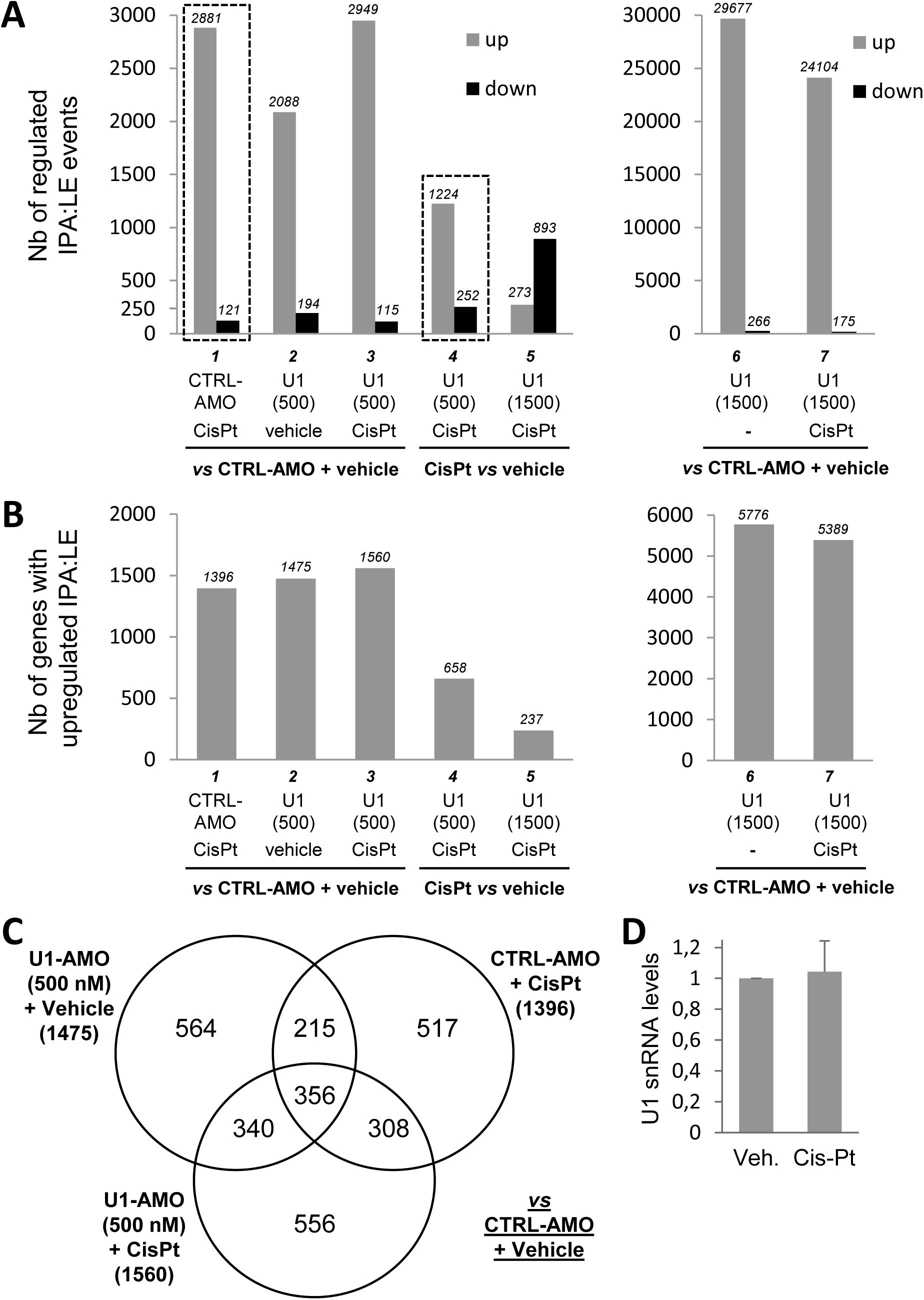
U1 snRNA blockade partially prevents cisplatin effect on the IPA:LE isoform ratio. A549 cells were electroporated with CTRL– and/or U1-AMO (with a total of 1500 nM AMO in every condition) and subsequently treated with either cisplatin at 5 µM or vehicle. 3’-seq analysis of IPA:LE isoform ratio regulation between the indicated conditions. U1, U1-AMO. **A**, Number of events (IPA sites) with up– or down-regulation of the IPA:LE isoform ratio. **B**, Number of genes with upregulated IPA:LE ratio. U1-AMO was used at 500 nM. **C**, Venn diagram of genes with upregulated IPA:LE ratio in different conditions when compared to control cells. **D**, RT-qPCR analysis of U1 snRNA levels in H358 cells treated with cisplatin or vehicle.

Firstly, the comparison of genes with IPA:LE events upregulated by moderate doses of either U1-AMO (500 nM) or cisplatin (5 µM) alone revealed a limited overlap of 39% (Fig. 5C), suggesting that these agents have at least in part different specificity towards IPA regulation. Consistently with this observation, cisplatin treatment did not appear to regulate U1 snRNA levels (Fig. 5D), which is in contrast with its downregulation upon ultraviolet-C irradiation (Devany et al. 2016).

Secondly, the combined treatment with U1-AMO at 500 nM and cisplatin upregulated about the same number of IPA:LE events (2949 events in 1560 genes) as each agent alone (Fig. 5A-B, lanes 1-3; Supplemental Fig. S1C and Supplemental Table S5). In addition, 64% of the genes with IPA:LE events upregulated by the combined agents were also regulated by at least one agent alone (Fig. 5C). Thus, the effects of U1-AMO and cisplatin on IPA:LE isoforms are less than additive on a global scale. (Nevertheless, U1-AMO and cisplatin may cooperate to regulate specific IPA:LE events, because we found 556 events that seem to be upregulated only by the combined agents (Fig. 5C)).

Thirdly, there were 2.4 times less (1224 *versus* 2881) IPA:LE events upregulated by cisplatin relative to vehicle if cells were transfected beforehand with 500 nM U1-AMO instead of CTRL-AMO (Fig. 5A, comparing lanes 1 and 4, and Supplemental Fig. S1E-F). Conversely, there were 2.1 times more downregulated events (252 *versus* 121). These data suggest that U1-AMO prevents a large subset of cisplatin effects on IPA:LE events.

## DISCUSSION

A reciprocal crosstalk between IPA and the DDR has emerged (see introduction). The U1 snRNA promotes transcription processivity in long genes by repressing IPA (Oh et al. 2017) and was involved in IPA regulation by ultraviolet-C irradiation (Devany et al. 2016), but little is known about its effects on DNA damage and on cell sensitivity to genotoxic and/or anticancer agents. The present study shows that U1 snRNA blockade decreases full-length transcripts in DNA repair genes, increases cisplatin-induced DNA damage, and sensitizes lung cancer cells to cisplatin. We also find that U1 snRNA blockade partially prevents cisplatin effect on the IPA:LE isoform ratio on a genome-wide scale (although this may not pertain to cisplatin sensitization).

Our finding that U1 snRNA blockade can sensitize lung cancer cells to cisplatin (Fig. 1) is consistent with the previous observation that U1 snRNA overexpression prevents apoptosis induction by ultraviolet-C irradiation, but the underlying mechanism was unknown (Devany et al. 2016). Our data show that U1 snRNA blockade increases cisplatin-induced DNA damage (Fig. 3), presumably due to the repression of multiple genes involved in the NER and FA pathways (Fig. 4), that play key roles in the repair of DNA crosslinks induced by this drug (Rocha et al. 2018). Several aspects would need to be further studied. First, we cannot exclude that U1-AMO effects on splicing (independently of IPA) may also contribute to its effects on cisplatin response, although such splicing effects of U1-AMO are less pronounced than its IPA effects (Kaida et al. 2010; Berg et al. 2012).

Second, while our data show that U1-AMO represses genes of the NER and FA pathways involved in the repair of DNA crosslinks, previous studies showed that CDK12 inhibition represses genes involved in the homologous recombination pathway, that is particularly important for the repair of double-strand breaks and at stalled replication forks (Chirackal Manavalan et al. 2019; Dubbury et al. 2018; Fan et al. 2020; Krajewska et al. 2019; Quereda et al. 2019; Tien et al. 2017). Thus, in the future, it would be interesting to compare the effects of targeting either U1 snRNA, CDK12 or both, on the various pathways of DNA repair and DDR signaling, and on cancer cell sensitivity to various genotoxic anticancer agents that induce different types of DNA lesions. This could help design novel drug combinations for anticancer therapy.

Third, our finding that in the absence of genotoxic agent, U1-AMO alone represses various cell cycle genes (Fig. 4B) and indeed impacts the cell cycle (Fig. 2) may be relevant to its anti-proliferative effects (Fig. 1A and 1C). However, the fact that U1-AMO partially prevents the cell cycle effects of cisplatin may limitate its ability to enhance cell growth inhibition by the drug, thus explaining the moderate sensitization we observed (Fig. 1B and 1D). This ought to be optimized and compared with CDK12 inhibitors.

Our data also suggest differences in the way cisplatin and ultraviolet-C treatment regulate the IPA:LE isoform ratio, with respect to the role of the U1 snRNA in this regulation. Indeed, a previous study showed that ultraviolet-C irradiation leads to a decrease of U1 snRNA levels and that its overexpression prevents IPA:LE isoform up-regulation in tested genes (Devany et al. 2016). In contrast, in the case of cisplatin treatment, U1 snRNA levels do not seem to be regulated (Fig. 5D) and U1 snRNA blockade partially prevents cisplatin effect on the IPA:LE isoform ratio on a genome-wide scale (Fig. 5A). Nevertheless, for both agents, the up-regulation of the IPA:LE isoform ratio is due at least in part to a decrease of transcription processivity, which decreases LE isoforms (Williamson et al. 2017; Devaux et al. 2025).

Altogether, this study increases our understanding of IPA isoform regulation by genotoxic agents and of DDR gene regulation at the IPA level, and identifies the U1 snRNA as a relevant target for cancer cell sensitization to genotoxic chemotherapy. In the future, building on these findings could help design novel anticancer therapeutic approaches.

## MATERIALS AND METHODS

### Cell culture, electroporation, and treatment

Cells were cultured in RPMI-1640-Glutamax (H358) or DMEM (A549, A549-R) medium (GibcoBRL, Life Technologies, Cergy Pontoise, France) supplemented with 10% (v/v) heat-inactivated fetal calf serum (GibcoBRL), in 5% CO_2_ at 37°C. Cells were electroporated using Cell Line Nucleofector™ Kit T (VCA-1002, Lonza). Following manufacturer’s recommendations, 1 million cells were mixed with 100 µL of buffer T kit with Supplement 1. The mix was put into the electroporation cuve provided by Lonza, with the indicated amounts of the U1-AMO (GGTATCTCCCCTGCCAGGTAAGTAT) and/or Standard Control morpholino oligonucleotides (Ctrl-AMO, Gene Tools, Philomath, Oregon, USA). The cuve was inserted into NucleofectorTM II and the A-023 (H358) or X-001 (A549/A549-R) program. For H358 or A549/A549-R respectively, 300 000 or 100 000 cells/mL were plated into 96-well plates, at 300 000 or 200 000 cells/mL for 12 well plates, and at 250 000 cells/mL for 6-well plates. For H358 or A549/A549-R respectively, 24 or 8 hrs after electroporation, cells were treated with either cisplatin (Selleckchem, Euromedex, Souffelweyersheim, France) or vehicle (DMSO) for 48 or 24 hrs.

### WST1 assay and FACS analysis

For cell growth analysis, cells were seeded in 96-well plates. Following cisplatin treatment, cell viability was assayed in triplicate wells using WST1 (Sigma Aldrich, France) according to the manufacturer’s instructions. For FACS analysis, cells were treated 2 hrs with BrdU 20 µM in a 6-well plate at the end of the indicated treatment. Cells were harvested then fixed with 70% cold ethanol and put at –20°C for storage. H358 and A549 cells were processed using FITC-BrdU Flow Kits according to the manufacturer’s instructions (BD Biosciences, San Jose, CA, USA cat.51–2354AK). Flow cytometric analysis of 10000 cells was performed on a FACScan flow cytometer (BD Biosciences) and data were recovered using the CellQuest software (BD Biosciences).

### Immunofluorescence analysis

Transfected cells grown on Marienfeld Superior cover glasses were wased twice with ice cold PBS, fixed using 4% paraformaldehyde in PBS during 20 min, washed three times with PBS, permeabilized for 10 min with PBS-0.1% Triton-X, washed twice with PBS, blocked for 5 min with PBS-5% BSA, incubated with Anti-γH2AX (phospho S139) rabbit antibody (ab26350, Abcam, 1:2000) at RT for 1 hour, washed twice with PBS, blocked again, incubated with F(ab’)2-rabbit anti-mouse IgG (H+L) cross-adsorbed secondary antibody, Alexa Fluor 488 (A-21206, Thermofischer, 1:1000) in PBS-5% BSA for 1 hour, washed twice with PBS, incubated at RT with PBS containing 4’,6-diamidino-2-phenylindole (DAPI) 0.1 µg/mL, and washed twice with PBS. Cover glasses were mounted on slides in PBS, glycerol 15%, 1.4-diazabicyclo-(2.2.2) octane (DABCO, Sigma) 100 mg/ml. For microscopy, acquisition was done with an exposure time of 50 ms for Trans-DIC, 20 ms for DAPI 405 and 50 ms for FITC. Z stacking was used for imaging DAPI and FITC with 31 panels for a range of ±5 µm around the acquisition point.

### RNA extraction

RNA from whole cells were extracted with TRIzol Reagent (Thermo Fisher Scientific) according to the manufacturer’s instructions, and 1 µl of GlycoBlue (Thermo Fisher Scientific) was added for RNA precipitation. RNA was treated with DNase I (TURBO DNA-free, Thermo Fisher Scientific). RNA samples were quantified using a Nanodrop 2000 spectrophotometer. For sequencing, RNA samples were analyzed using an RNA 2100 Bioanalyzer (Agilent).

### RT-qPCR

Reverse transcription was performed on RNA using SuperScript III Reverse Transcriptase (ThermoFisher Scientific) and oligo-dT primers (except for U1 snRNA, for which random primers were used). Quantitative PCR (qPCR) was performed using Power SYBR Green PCR Master Mix (ThermoFisher Scientific) on a CFX96 Real-Time PCR Detection System (BioRad). Primer sequences are given in Supplemental Table S6.

### 3’-seq experiments and bioinformatic analysis

All the 3’-seq analyses in this study were carried out in parallel and on 4 biological replicates. Each replicate experiment included 6 conditions: CTRL-AMO or U1-AMO at different doses, with or without cisplatin. 3’-seq libraries were prepared with QuantSeq 3’ mRNA-Seq Library Prep Kit REV for Illumina (Lexogen) using 500 ng of DNase I treated RNA (n=4 for each condition) following manufacturer’s instructions. Purified libraries were quantified with Quant-iT Picogreen dsDNA kit (Thermo Fisher Scientific) and run on Experion automated electrophoresis system (BIO-RAD). Pooled libraries were quantitated by qPCR (KAPA Library Quantification Kits Illumina Platforms, Roche), diluted to 12 pM, and subjected to single-end, 50 bp sequencing using the NovaSeq 6000 machine (Illumina).

For each sample, raw reads were trimmed to remove uninformative nucleotides due to primer sequences. Trimmed reads of 25 bp or more were aligned on the human reference genome (hg19) using Bowtie 2 (version 2.2.5) (Langmead and Salzberg 2012). Only reads with a mapping quality score (MAPQ) of 20 or more were retained (SAMtools version 1.1) for downstream analysis (Li et al. 2009). Reads were then clustered along the genome using BEDTools (version 2.17.0) (Quinlan and Hall 2010), allowing a maximum distance of 50 bp and a minimum number of 5 reads per peak. Peaks with a stretch of 6 consecutive As (or 8 As out of 9 nucleotides) within 50 bp downstream were filtered out, as they are likely due to internal priming of Oligo(dT). Overlapping peaks from all samples of the compared conditions were merged to define a common set of genomic windows corresponding to poly(A) sites. To annotate peak location within genes, gene coordinates were obtained on the basis of overlapping RefSeq transcripts with the same gene symbol. Peaks overlapping any intronic region of a gene were classified as intronic poly(A) (IPA) peaks. Peaks overlapping the last exon of a gene were classified as LE peaks. Differential analyses between two conditions were done using four independent biological replicates per condition.

To compare the regulation of each IPA to the regulation of the gene’s last exon (taken as the sum of the peaks in this exon), we used DESeq2 (version 1.4.5) (Love et al. 2014) and the following statistical model:

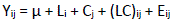

where Y_ij_ is the normalized counts of peak i in biological condition j, μ is the mean, L_i_ is the peak localization (IPA or LE), C_j_ is the biological condition, (LC)_ij_ is the interaction between peak localization and biological condition, and E_ij_ is the residual. P-values and adjusted P-values (Benjamini-Hochberg) were calculated. Data with p < 0.05 are shown. Among regulated genes, we identified those with at least 4-fold downregulation of the last exon. Functional gene annotation analyses were done using the DAVID software (Huang et al. 2009b, 2009a), using the human genome as a reference.

### Statistical analyses

For each experimental analysis, at least three independent experiments were performed. In all bar charts, error bars represent the standard error of the mean (SEM) that is the standard deviation divided by the square root of sample number. A Student’s t-test was used, except in Fig. 3, where a Welch t-test was used. Tests were considered significant if p < 0.05.

## DATA AVAILABILITY

The datasets generated in this study were deposited in the Gene Expression Omnibus repository (GSE342386, private token: whclcmukbxwzrqz). The complete bioinformatics pipeline for 3’-seq analysis of IPA (3’-SMART package) can be freely downloaded at GitHub (https://github.com/bioinfo-pf-curie/3-SMART) and can be run through a configuration file and a simple command line.

## SUPPLEMENTAL MATERIAL

Supplemental Material is available for this article.

## ACKNOWLEDGMENTS

This work was supported by grants of Institut Curie (ICGex) to MD and of Ligue Nationale Contre le Cancer (équipe labellisée) to SV, and by fellowships from Ministère de l’Enseignement Supérieur et de la Recherche, Université Paris Saclay and Institut Curie to AD. We thank the CurieCoreTech – Cytometry (CYTPIC) platform and the Light Microscopy facility (Multimodal Imaging Center) of Institut Curie. The Next Generation Sequencing platform (ICGex) of Institut Curie is supported by grants ANR-10-EQPX-03 (Equipex) and ANR-10-INBS-09-08 (France Génomique Consortium) from the Agence Nationale de la Recherche (“Investissements d’Avenir” program), by the Cancéropôle Ile-de-France and by the SiRIC-Curie program – SiRIC Grant INCa-DGOS-4654. We thank Simon Coiba for help in RT-qPCR analyses.

## AUTHORS’ CONTRIBUTIONS

Experiments were performed by AD under the supervision of MD. Bioinformatics analyses were performed by CML under the supervision of MD. MD wrote the paper with help of AD and SV.

## REFERENCES

1. Andersen PK, Lykke-Andersen S, Jensen TH. 2012. Promoter-proximal polyadenylation sites reduce transcription activity. Genes Dev 26: 2169–2179.

2. Ashrafi A, Akter Z, Modareszadeh P, Modareszadeh P, Berisha E, Alemi PS, Chacon Castro MDC, Deese AR, Zhang L. 2022. Current Landscape of Therapeutic Resistance in Lung Cancer and Promising Strategies to Overcome Resistance. Cancers (Basel) 14: 4562.

3. Bartkowiak B, Yan CM, Soderblom EJ, Greenleaf AL. 2019. CDK12 Activity-Dependent Phosphorylation Events in Human Cells. Biomolecules 9: 634.

4. Berg MG, Singh LN, Younis I, Liu Q, Pinto AM, Kaida D, Zhang Z, Cho S, Sherrill-Mix S, Wan L, et al. 2012. U1 snRNP determines mRNA length and regulates isoform expression. Cell 150: 53–64.

5. Chakraborty A, Cadix M, Relier S, Taricco N, Alaeitabar T, Devaux A, Labbé CM, Martineau S, Heneman-Masurel A, Gestraud P, et al. 2022. Compartment-specific and ELAVL1-coordinated regulation of intronic polyadenylation isoforms by doxorubicin. Genome Res 32: 1271–1284.

6. Chirackal Manavalan AP, Pilarova K, Kluge M, Bartholomeeusen K, Rajecky M, Oppelt J, Khirsariya P, Paruch K, Krejci L, Friedel CC, et al. 2019. CDK12 controls G1/S progression by regulating RNAPII processivity at core DNA replication genes. EMBO Rep 20: e47592.

7. Choi SH, Martinez TF, Kim S, Donaldson C, Shokhirev MN, Saghatelian A, Jones KA. 2019. CDK12 phosphorylates 4E-BP1 to enable mTORC1-dependent translation and mitotic genome stability. Genes Dev 33: 418–435.

8. Ciccia A, Elledge SJ. 2010. The DNA damage response: making it safe to play with knives. Mol Cell 40: 179–204.

9. Damgaard CK, Kahns S, Lykke-Andersen S, Nielsen AL, Jensen TH, Kjems J. 2008. A 5’ splice site enhances the recruitment of basal transcription initiation factors in vivo. Mol Cell 29: 271– 278.

10. Devany E, Park JY, Murphy MR, Zakusilo G, Baquero J, Zhang X, Hoque M, Tian B, Kleiman FE. 2016. Intronic cleavage and polyadenylation regulates gene expression during DNA damage response through U1 snRNA. Cell Discov 2: 16013.

11. Devaux A, Tanaka I, Fouilleul Q, Heneman-Masurel A, Cadix M, Michallet S, Chakraborty A, Labbé CM, Fontrodona N, Sahoo S, et al. 2025. Identification of microprotein-coding intronic polyadenylation isoforms and function in genotoxic anticancer drug response. Genome Biol 26: 366.

12. Dubbury SJ, Boutz PL, Sharp PA. 2018. CDK12 regulates DNA repair genes by suppressing intronic polyadenylation. Nature 564: 141–145.

13. Dutertre M, Chakrama FZ, Combe E, Desmet F-O, Mortada H, Polay Espinoza M, Gratadou L, Auboeuf D. 2014a. A recently evolved class of alternative 3’-terminal exons involved in cell cycle regulation by topoisomerase inhibitors. Nat Commun 5: 3395.

14. Dutertre M, Lambert S, Carreira A, Amor-Guéret M, Vagner S. 2014b. DNA damage: RNA-binding proteins protect from near and far. Trends Biochem Sci 39: 141–149.

15. Dutertre M, Sfaxi R, Vagner S. 2021. Reciprocal Links between Pre-messenger RNA 3’-End Processing and Genome Stability. Trends Biochem Sci 46: 579–594.

16. Fan Z, Devlin JR, Hogg SJ, Doyle MA, Harrison PF, Todorovski I, Cluse LA, Knight DA, Sandow JJ, Gregory G, et al. 2020. CDK13 cooperates with CDK12 to control global RNA polymerase II processivity. Sci Adv 6: eaaz5041.

17. Gunderson SI, Polycarpou-Schwarz M, Mattaj IW. 1998. U1 snRNP inhibits pre-mRNA polyadenylation through a direct interaction between U1 70K and poly(A) polymerase. Mol Cell 1: 255–264.

18. Huang DW, Sherman BT, Lempicki RA. 2009a. Bioinformatics enrichment tools: paths toward the comprehensive functional analysis of large gene lists. Nucleic Acids Res 37: 1–13.

19. Huang DW, Sherman BT, Lempicki RA. 2009b. Systematic and integrative analysis of large gene lists using DAVID bioinformatics resources. Nat Protoc 4: 44–57.

20. Kaida D, Berg MG, Younis I, Kasim M, Singh LN, Wan L, Dreyfuss G. 2010. U1 snRNP protects pre-mRNAs from premature cleavage and polyadenylation. Nature 468: 664–668.

21. Krajewska M, Dries R, Grassetti AV, Dust S, Gao Y, Huang H, Sharma B, Day DS, Kwiatkowski N, Pomaville M, et al. 2019. CDK12 loss in cancer cells affects DNA damage response genes through premature cleavage and polyadenylation. Nat Commun 10: 1757.

22. Kwek KY, Murphy S, Furger A, Thomas B, O’Gorman W, Kimura H, Proudfoot NJ, Akoulitchev A. 2002. U1 snRNA associates with TFIIH and regulates transcriptional initiation. Nat Struct Biol 9: 800–805.

23. Langmead B, Salzberg SL. 2012. Fast gapped-read alignment with Bowtie 2. Nat Methods 9: 357–359.

24. Li H, Handsaker B, Wysoker A, Fennell T, Ruan J, Homer N, Marth G, Abecasis G, Durbin R, 1000 Genome Project Data Processing Subgroup. 2009. The Sequence Alignment/Map format and SAMtools. Bioinformatics 25: 2078–2079.

25. Love MI, Huber W, Anders S. 2014. Moderated estimation of fold change and dispersion for RNA-seq data with DESeq2. Genome Biol 15: 550.

26. Michels J, Vitale I, Galluzzi L, Adam J, Olaussen KA, Kepp O, Senovilla L, Talhaoui I, Guegan J, Enot DP, et al. 2013. Cisplatin resistance associated with PARP hyperactivation. Cancer Res 73: 2271– 2280.

27. Mimoso CA, Adelman K. 2023. U1 snRNP increases RNA Pol II elongation rate to enable synthesis of long genes. Mol Cell 83: 1264–1279.e10.

28. Oh J-M, Di C, Venters CC, Guo J, Arai C, So BR, Pinto AM, Zhang Z, Wan L, Younis I, et al. 2017. U1 snRNP telescripting regulates a size-function-stratified human genome. Nat Struct Mol Biol 24: 993–999.

29. Quereda V, Bayle S, Vena F, Frydman SM, Monastyrskyi A, Roush WR, Duckett DR. 2019. Therapeutic Targeting of CDK12/CDK13 in Triple-Negative Breast Cancer. Cancer Cell 36: 545–558.e7.

30. Quinlan AR, Hall IM. 2010. BEDTools: a flexible suite of utilities for comparing genomic features. Bioinformatics 26: 841–842.

31. Rocha CRR, Silva MM, Quinet A, Cabral-Neto JB, Menck CFM. 2018. DNA repair pathways and cisplatin resistance: an intimate relationship. Clinics (Sao Paulo) 73: e478s.

32. Tien JF, Mazloomian A, Cheng S-WG, Hughes CS, Chow CCT, Canapi LT, Oloumi A, Trigo-Gonzalez G, Bashashati A, Xu J, et al. 2017. CDK12 regulates alternative last exon mRNA splicing and promotes breast cancer cell invasion. Nucleic Acids Res 45: 6698–6716.

33. Vagner S, Rüegsegger U, Gunderson SI, Keller W, Mattaj IW. 2000. Position-dependent inhibition of the cleavage step of pre-mRNA 3’-end processing by U1 snRNP. RNA 6: 178–188.

34. Vorlová S, Rocco G, Lefave CV, Jodelka FM, Hess K, Hastings ML, Henke E, Cartegni L. 2011. Induction of antagonistic soluble decoy receptor tyrosine kinases by intronic polyA activation. Mol Cell 43: 927–939.

35. Williamson L, Saponaro M, Boeing S, East P, Mitter R, Kantidakis T, Kelly GP, Lobley A, Walker J, Spencer-Dene B, et al. 2017. UV Irradiation Induces a Non-coding RNA that Functionally Opposes the Protein Encoded by the Same Gene. Cell 168: 843–855.e13.

