## Supplementary material for "U1 snRNA blockade regulates DNA repair genes, DNA damage, and cisplatin sensitivity of lung cancer cells": Suppl Figures

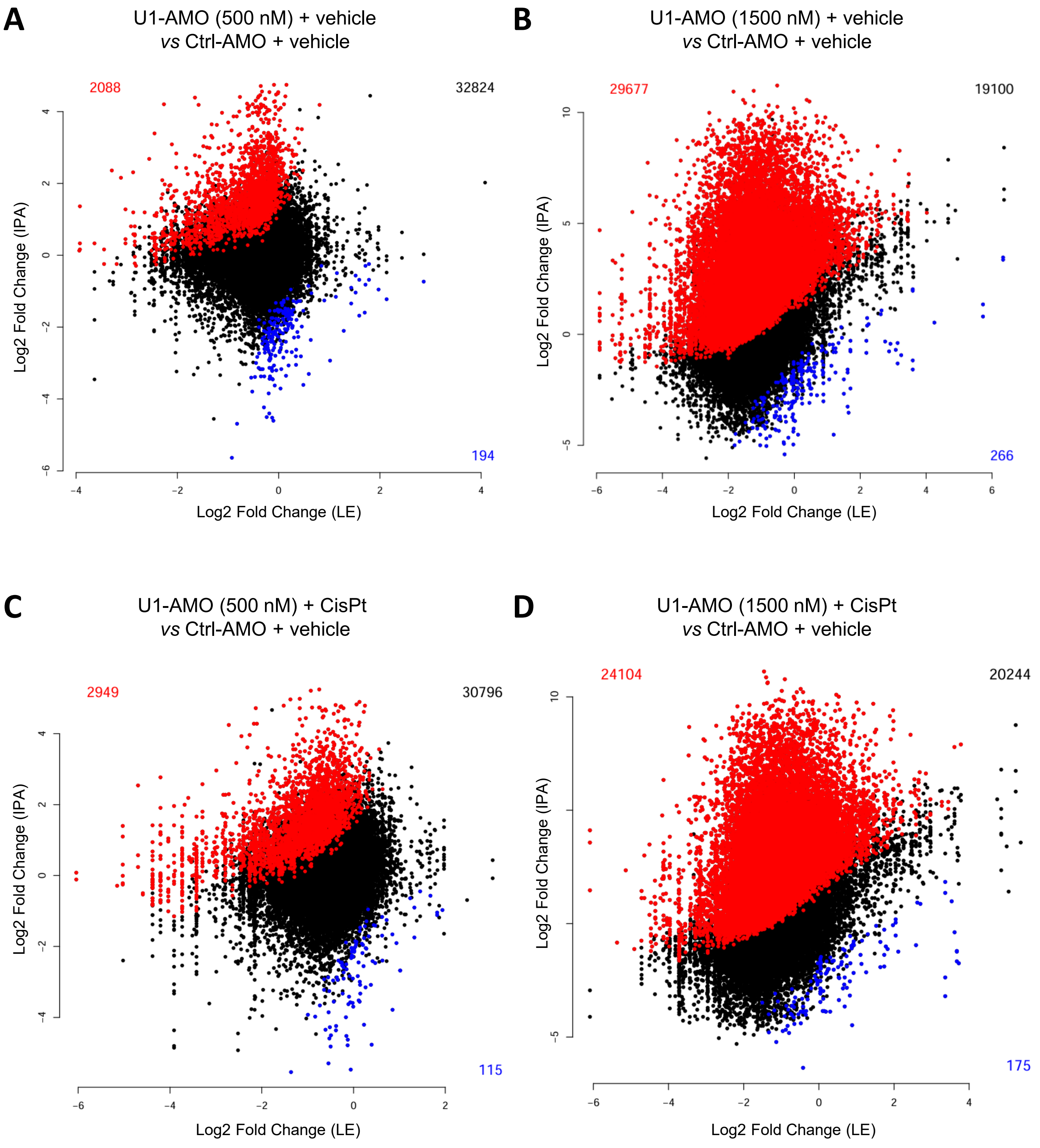


**Supplementary Figure S1** (to be continued on next page)

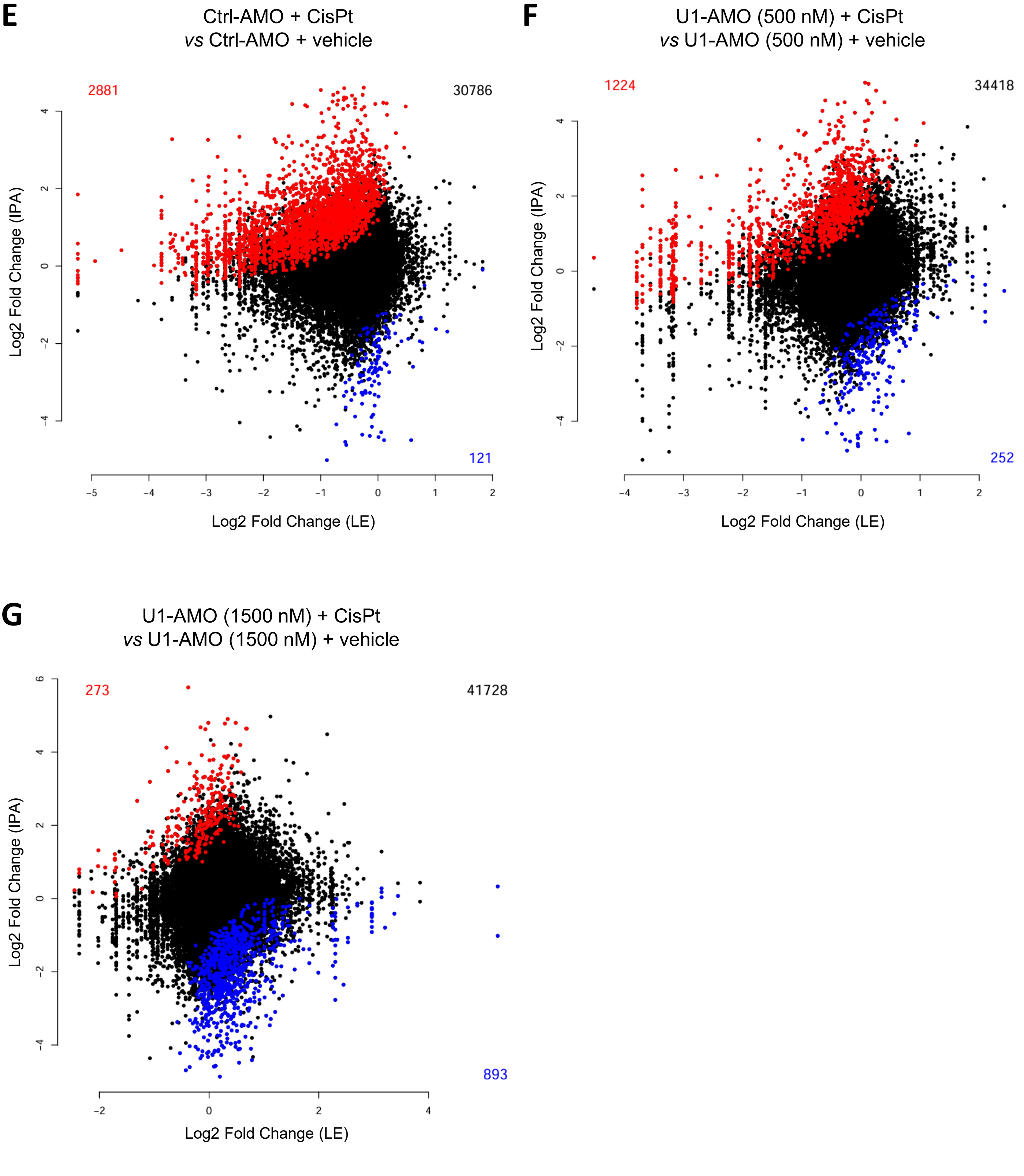


**Supplementary Figure S1:** Scatter plots corresponding to the 3'-seq analyses of IPA and last-exon (LE) isoforms regulation between the indicated conditions. Each dot corresponds to an IPA site. Red and blue dots, IPA sites with significantly up- or down-regulated IPA:LE isoform ratio, respectively.
